# ADAR RNA editing for cardiovascular disease: Targeting *B4GALT1* to modulate lipid metabolism through reduced galactosyltransferase activity

**DOI:** 10.1101/2025.10.30.685343

**Authors:** Francesco De Chiara, Eva Coll-de la Rubia, Petra de Bruijn, Galja Pletikapic, Nicole Cnubben, Lianne Stevens, Emy Theunissen, Janne J. Turunen, Anita M. van den Hoek, Hans M. G. Princen, Wouter Beumer, Seda Yilmaz-Elis, Bart Klein, Sjef de Kimpe, Gerard Platenburg

## Abstract

Cardiovascular disease remains a leading cause of mortality despite current therapies targeting low-density lipoprotein cholesterol (LDL-C). Beta-1,4-galactosyltransferase 1 (B4GALT1), a central glycosyltransferase enzyme that regulates lipoprotein metabolism and hemostasis, is a promising therapeutic target. To evaluate its therapeutic potential as a protective variant, *B4galt1* p.Asn352Ser was introduced into the ribonucleic acid (RNA) of healthy wild-type and APOE*3-Leiden.CETP transgenic mice, a well-established model for hyperlipidemia with a humanized lipoprotein metabolism, using editing oligonucleotides and the endogenous adenosine deaminases acting on RNA enzymes. The impact on hepatic glycosylation and systemic lipid homeostasis was investigated using multi-omics profiling. Editing of *B4galt1* messenger RNA (∼9−18%) resulted in substantial reductions in total cholesterol (–61%), apolipoprotein B (–72%), LDL-C (–30%), fibrinogen (–55%) (all *p* < 0.05), and triglycerides (–28%; *p* > 0.05), without altering *B4galt1* expression. Proteomics of plasma and liver identified early suppression of lipogenesis and lipoprotein assembly, followed by sustained suppression of cholesterol biosynthesis and coagulation pathways. Glycomic analysis revealed remodeling of circulating glycoprotein architecture, consistent with altered B4GALT1 activity. Transcript–protein concordance was strongest in lipid pathways, while glycosylation and coagulation showed domain-specific regulatory patterns. These findings support targeting *B4GALT1* using RNA editing to reduce cardiometabolic risk.

## Introduction

Despite the availability of effective treatments, patients with cardiovascular disease (CVD) remain at a high risk of adverse events including increased mortality.^1, 2^ This persistent burden underscores the need for therapies that can precisely modulate the metabolic processes driving CVD. Protective variants of naturally occurring circulating enzymes offer a compelling model for targeted metabolic interventions. A notable example is the p.Asn352Ser variant of beta-1,4-galactosyltransferase 1 (*B4GALT1*), which is enriched in the US Amish community and is associated with lower low-density lipoprotein cholesterol (LDL-C) and fibrinogen levels, as well as reduced cardiovascular risk, compared with the US population overall.^2^ Genetic studies of populations carrying the protective *B4GALT1* variant demonstrate how even subtle changes at conserved catalytic or substrate-binding residues can rewire enzymatic activity, leading to clinically meaningful metabolic protection.^2, 3^ These findings highlight the therapeutic potential of targeted modulation of enzyme activity to mimic the protective effects of naturally occurring genetic variants.

B4GALT1 has emerged as a promising therapeutic target for CVD, owing to its central role in glycosylation and the subsequent modulation of lipoprotein metabolism, coagulation, and clearance.^4^ This enzyme catalyzes the terminal β1,4-galactosylation of N-linked glycans on key secretory proteins such as apolipoprotein B (ApoB), LDL, and fibrinogen, influencing the functional stability and rate of clearance from circulation.^5–7^ The activity of B4GALT1 depends on uridine diphosphate (UDP)-galactose, which is regenerated by UDP-galactose-4-epimerase (GALE), placing GALE upstream in the same metabolic pathway.^8, 9^ Disruption of glycosylation impairs receptor recognition and clearance, promoting atherogenesis.^5–7^ The glycosylation of circulating proteins, including lipoproteins and coagulation factors, is a fundamental yet underappreciated aspect of cardiometabolic regulation.^6, 7^

B4GALT1 is broadly expressed across tissues, consistent with its essential role in N-glycan maturation.^10^ In the context of cardiometabolic biology, its hepatic activity is particularly relevant, as hepatocytes produce circulating ApoB-containing lipoproteins and fibrinogen.^11^ The protective human p.Asn352Ser variant in B4GALT1 is associated with lower circulating LDL-C and fibrinogen levels, and has been shown to reduce galactosyltransferase activity by approximately 50%, leading to altered glycosylation of circulating proteins.^2^ This partial loss of enzymatic activity, while preserving overall protein expression, supports therapeutic modulation of B4GALT1 rather than complete inhibition.

Leveraging ribonucleic acid (RNA) editing to introduce a protective B4GALT1 variant represents a novel therapeutic strategy to enhance glycosylation-dependent clearance mechanisms and reduce cardiovascular risk. Axiomer^TM^ technology is an RNA editing platform that employs chemically modified editing oligonucleotides (EONs) designed to pair with a target messenger RNA (mRNA) to recruit endogenous adenosine deaminase acting on RNA (ADAR), facilitating site-specific A-to-I editing.^12^ This approach holds substantial therapeutic potential, as it can correct numerous disease-related mutations, modulate protein functions, or introduce protective variants.^12^ RNA editing of enzymatic regulators can reshape post-translational processes such as glycosylation and proteolysis, influencing signaling and protein fate decisions.^12^ By leveraging endogenous ADAR enzymes, Axiomer^TM^ avoids the need for potentially immunogenic viral vectors, offering a scalable and reversible alternative to conventional gene editing methods.^13^ The high specificity of this platform reduces the risk of unintended consequences, and its flexible design enables the targeting of different RNA species across multiple tissues.^13^

RNA editing represents a paradigm shift in therapeutic development, enabling the introduction of evolutionary advantageous variants into individuals who do not naturally carry them, while preserving natural gene regulation. In contrast to gene knockdown strategies such as clustered regularly interspaced short palindromic repeats (CRISPR) or small interfering RNA (siRNA), which disrupt or silence genes, RNA editing offers precise and tunable modulation of enzyme activity.^13^ Here, we mechanistically dissect the functional consequences of introducing the protective p.Asn352Ser variant into hepatic *B4galt1* transcripts, focusing on downstream alterations in glycoprotein composition, metabolic flux, and pathway-level regulation in wild-type and hyperlipidemic mice. Specifically, we describe two EONs (EON-1 and EON-2) developed through an iterative refinement process, with each successive generation incorporating structural and chemical refinements based on prior experimental insights. Using the most advanced candidates from each generation, we characterized the downstream molecular responses driving the observed improvements in clinical biomarkers. Through integrative transcriptomic and proteomic analyses, we highlight RNA editing as a mechanistically precise approach to reprogram metabolic networks by modulating nodal enzymatic activity, offering a targeted strategy to address residual cardiometabolic risk.

## Results

To evaluate the therapeutic potential of the B4GALT1 p.Asn352Ser variant, we employed a sequential translation-focused framework. Because the target editing site is conserved between species, we utilized EONs engineered to be compatible with both human and murine *B4GALT1/B4galt1* transcripts. The study progressed through three critical phases: (1) *in vivo* proof-of-concept, using a first-generation lead (EON-1) in wild-type C57BL/6J mice to establish pharmacokinetic/pharmacodynamic correlations and hepatic editing efficiency; (2) *in vitro* refinement, using primary human hepatocytes (PHH) to optimize structure-activity relationships for human target engagement; and (3) disease-model validation, where an optimized second-generation lead (EON-2) was administered chronically to Western diet-fed APOE*3-Leiden.CETP (E3L.CETP) mice. This final stage was designed to assess the impact of sustained *B4galt1* mRNA editing on complex lipid metabolism, coagulation, galactosyltransferase activity, and the systemic multi-omics landscape under conditions of metabolic stress.

### B4galt1 mRNA editing induces transient molecular remodeling and early metabolic benefits

In the *in vivo* proof-of-concept study, a single 2 mg/kg dose of lipid nanoparticles (LNP)-formulated EON-1 was administered to wild-type C57BL/6J mice. Plasma concentration peaked rapidly at approximately 10 minutes post dose, followed by rapid clearance (Figure 1A). Complementarily, hepatic accumulation of EON-1 peaked at 24 hours and remained detectable for up to 7 days, consistent with efficient liver uptake and retention (Figure 1A and 1B). This temporal lag phase is characteristic of LNP delivery systems in which the initial rapid distribution of the oligonucleotide cargo from the blood to the hepatic extracellular space is followed by a more gradual process of endocytosis and intracellular release. This pharmacokinetic profile closely aligned with the pharmacodynamic onset: *B4galt1* mRNA editing was first detectable at 24 hours, reaching approximately 6% at this peak and gradually declining thereafter, while remaining measurable up to 7 days post treatment (Figure 1B).

**Figure 1.**
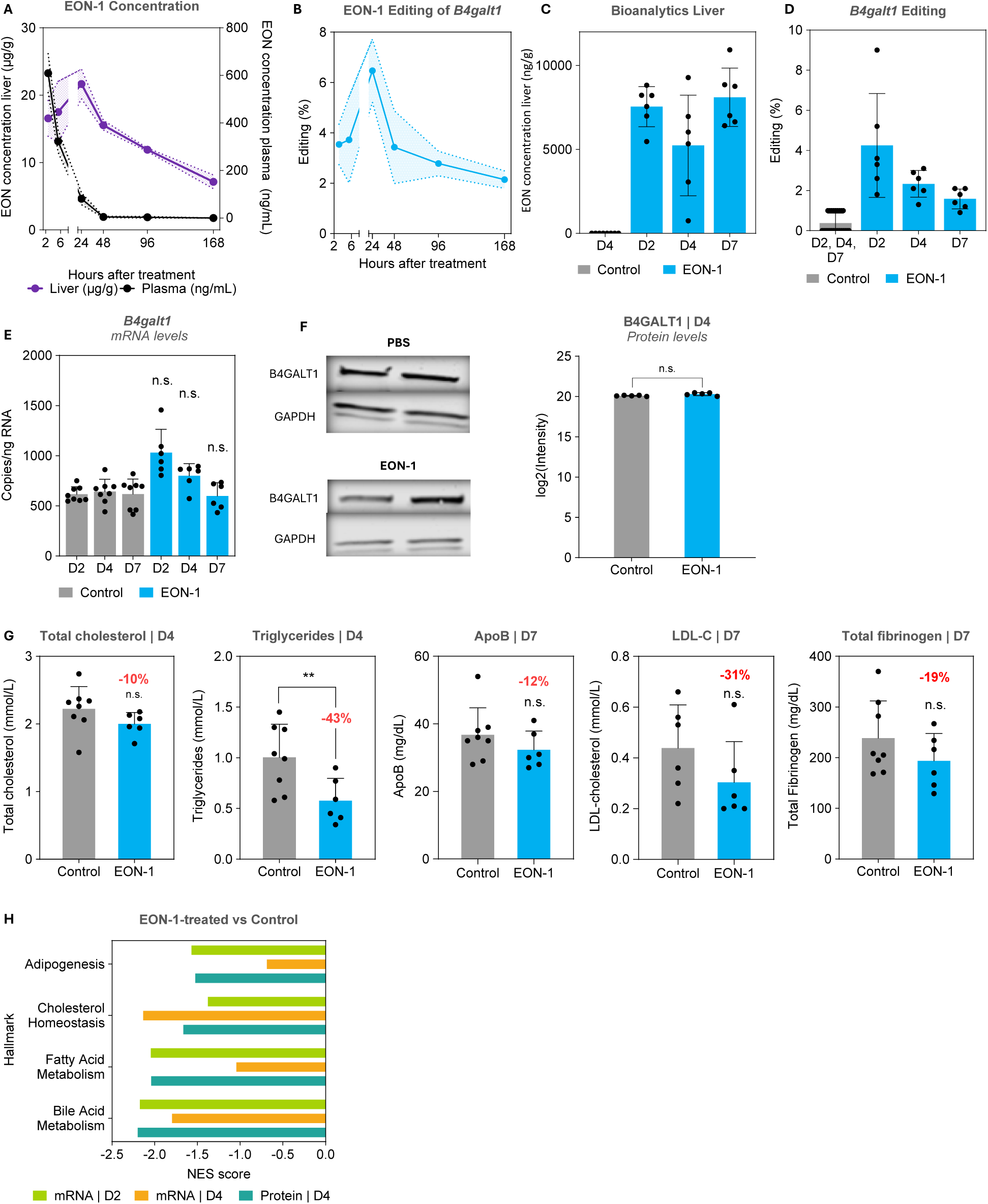
*B4galt1* editing induces transient molecular remodelling and early metabolic benefits. Plasma EON concentration was measured from 10 minutes to 168 hours, while EON liver concentration was assessed from 2 to 168 hours post treatment versus control: (A–B). Assessed at days 2, 4, and 7 post treatment versus control. (C) Hepatic accumulation of EON-1. (D) Mean levels of *B4galt1* mRNA editing. (E) Mean *B4galt1* mRNA expression levels (dPCR). (F) *B4galt1* protein abundance (western blot and quantitative proteomics). (G) Mean reductions in cholesterol, triglycerides, ApoB, and fibrinogen. (H) Editing-induced effect on adipogenesis, cholesterol homeostasis, fatty acid metabolism, and bile acid metabolism (NES score). Error bars represent standard deviation. For omics analyses, only pathways with adjusted *p* values (padj) < 0.05 were considered significant. \**p* □<□ 0.05 \*\**p* □<□ 0.01. ApoB, apolipoprotein B; BGALT1, beta-1,4-galactosyltransferase 1; dPCR, digital polymerase chain reaction; D4 / D7, day 4 / Day 7; EON, editing oligonucleotide; GAPDH, Glyceraldehyde 3-phosphate dehydrogenase; LDL, Low-density lipoprotein cholesterol; mRNA, messenger RNA; NES, normalized enrichment score; n.s. Not significant; PBS, phosphate-buffered saline.

In the follow-up pharmacodynamic exploratory study, *in vivo* administration of a single 3 mg/kg dose of EON-1 encapsulated in LNPs in wild-type C57BL/6J mice resulted in a similar EON liver concentration and editing percentage (Figure 1C and 1D). Importantly, no significant changes were detected in overall *B4galt1* mRNA expression levels (as determined by digital polymerase chain reaction [dPCR]) or protein abundance (assessed by western blot and semi-quantitative proteomics) at days 2, 4, or 7 post treatment (Figure 1E and 1F).

By day 4, EON-1 treatment led to significant reductions in triglycerides (−43%, *p* < 0.01), with trends toward reduction in total cholesterol (−10%) that did not reach statistical significance in this exploratory study. By day 7, further non-significant trends were observed for ApoB (−12%), LDL-C (−31%), and fibrinogen (−19%) (Figure 1G).

Transcriptomic and proteomic analyses revealed dynamic temporal regulation following *B4galt1* editing, with a pronounced initial surge in gene expression changes at day 2, followed by refinement of the transcriptional response by day 4 (Figure S1A, S1B). The editing-induced effect displayed a distinctive wave-like pattern, marked initially by transient suppression of bile acid metabolism, fatty acid metabolism, and adipogenesis-related genes on day 2, which recovered by day 4, as shown by a negative normalized enrichment score (NES) (Figure 1H). Conversely, cholesterol homeostasis remained persistently downregulated at both mRNA and protein levels (Figure 1H). This effect was primarily driven by reduced expression of *CYP8B1*, which encodes *sterol 12-alpha-hydroxylase*, an enzyme that regulates the ratio of cholic acid to chenodeoxycholic acid in the bile acid synthetic pathway (Figure S1C). In this figure, labeled genes represent those with statistically significant changes in abundance (adjusted *p*-value < 0.05).

Downregulation of *Gale* mRNA was also observed, without a corresponding reduction in protein levels (Figure S1C). This downregulation may reflect transcriptional adaptation to reduced galactosylation demand following *B4galt1* editing.^14, 15^ Given the generally longer half-life of metabolic enzymes relative to their transcripts, this likely represents a transient, buffered response, highlighting temporal and regulatory decoupling between transcriptomic and proteomic layers in galactose metabolism (Figure S1C).^16, 17^

### Iterative refinement of editing oligonucleotides reveals structure–function relationships driving RNA editing efficiency

Building on this initial proof-of-concept molecule (EON-1) to establish the feasibility of ADAR-mediated *B4galt1* editing in the mouse liver, we pursued an iterative refinement strategy based on sequence features such as length, symmetry, chemical modifications, and RNA thermodynamic properties to improve editing efficiency. To compare the pharmacological profiles of successive *B4GALT1*-targeting oligonucleotides (EON-1 and EON-2), we derived key dose–response parameters from *in vitro* transfection studies in PHH as *in vitro* refinement. EON-2 exhibited the highest E_max_ (14.2% vs EON-1, 5.1%) and the lowest EC₅₀ (21.9□ nM vs EON-1, 27.8□ nM), indicating superior efficacy and potency compared with EON-1. Area under the curve (AUC) analysis reinforced these trends, with EON-2 demonstrating substantially higher overall editing performance (AUC = 14.9) compared with EON-1 (5.0). Notably, EON-2 also showed a steeper Hill slope compared with EON-1, reflecting tighter dose-dependent control (Figure 2A).

**Figure 2.**
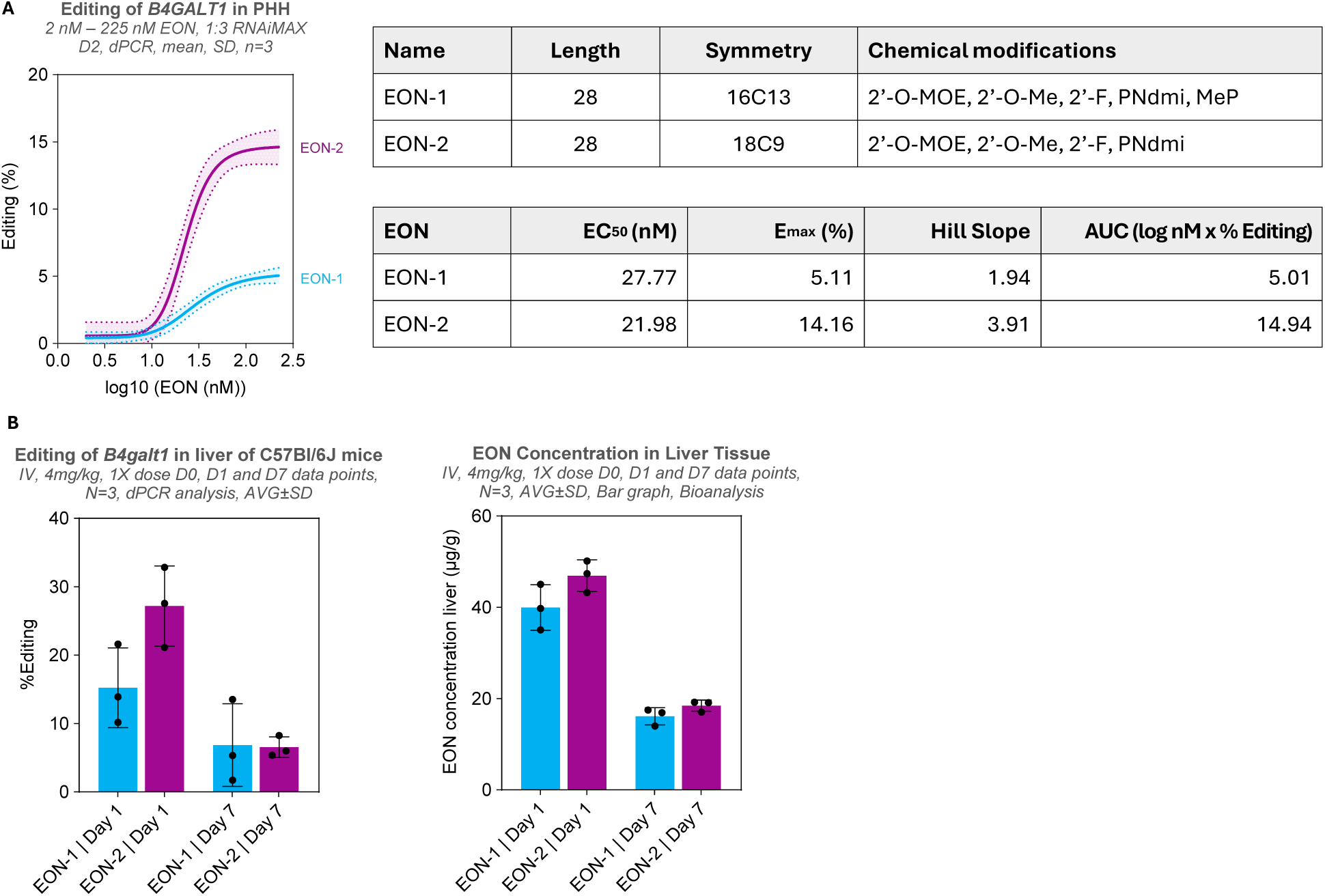
Iterative refinement of editing oligonucleotides reveals structure–function relationships driving RNA editing efficiency. Editing of *B4galt/B4GALT1* with EON-1 versus EON-2 in: (A) PHH at day 2, assessed using dPCR. (B) C57BL/6J mice over 7 days. AUC, area under the curve; BGALT1, beta-1,4-galactosyltransferase 1; dPCR, digital polymerase chain reaction; EC₅₀, half maximal effective concentration; Emax, maximum effect (maximum response); EON, editing oligonucleotide; 2′MOE, 2′-O-methoxyethyl; 2′F, 2′-fluoro; MeP, Methylphosphonate; PNdmi, dimethylamino-modified phosphoramidate; SD, standard deviation.

These functional differences map to specific structural and chemical features of the EONs. While both include 2’-O-methoxyethyl (2′-O-MOE), 2’-O-methylation (2′-O-Me), and 2’-fluoro (2′-F) sugar modifications, they differ in their internucleoside linkage composition. Additionally, factors such as the symmetry around the nucleotide opposite the target adenosine in the target sequence may also influence activity. Taken together, these findings show how, through systematic variation of symmetry around the editing site (16C13 to 18C9), mismatched nucleotides at positions +8 to +11 relative to the editing site and backbone chemistry, we identified EON-2 as having superior editing characteristics (Figure 2A), enabling both precise enzyme modulation and durable metabolic reprogramming.

Following the *in vitro* evaluations, we performed a pilot *in vivo* confirmatory study in C57BL/6J mice to verify that the potency hierarchy observed in PHH translated to mice. Mirroring the *in vitro* results, EON-2 out-performed EON-1 *in vivo*, achieving ∼27% *B4galt1* mRNA editing compared with ∼15% for EON-1 at day 1, despite indistinguishable hepatic oligonucleotide levels at the same time point (Figure 2B). Notably, although the EONs were engineered against the human *B4GALT1* transcript sequence and contain one additional nucleotide mismatch relative to the murine transcript, robust editing was achieved, indicating tolerance of this mismatch at the target site. By day 7, editing efficiency for both EONs had converged to ∼10%, coinciding with a parallel decline of 60% (*p*-value < 0.01) in liver EON concentrations (from 43.2 ± 5.4 to 17.3 ± 1.9 µg/g⁻¹) at day 1 and day 7, respectively (Figure 2B). These data confirm that the optimized chemistry of EON-2 translates to superior *in vivo* potency without altering pharmacokinetic behavior, thereby validating the feasibility of redirecting RNA editing in mice using an oligonucleotide engineered for the human *B4GALT1* transcript sequence. EON-2 emerged as the EON that satisfied editing potency both *in vitro* and *in vivo*, as well as manufacturability criteria, and was therefore advanced to the initial *in vivo* validation study.

### Enhanced B4galt1 RNA editing efficiency achieves sustained and improved lipid modulation and optimized therapeutic outcomes in hyperlipidemic, Western diet-fed E3L.CETP mice

In the disease-model validation study, EON-2 demonstrated improvement in editing efficiency, achieving nearly 15% hepatic *B4galt1* mRNA editing as early as day 1 in Western diet–fed E3L.CETP mice, in which dietary challenge induces hyperlipidemia. The E3L.CETP model is a well-established and clinically relevant model for metabolic and CVDs that is responsive to all lipid-lowering interventions that are being used in the clinic, including statins, fibrates, ezetimibe, and PCSK9 and ANGPTL3-inhibitors.^18–21^ Robust editing levels persisted until day 45, confirming consistent and durable RNA modification (Figure 3B, left panel). Bioanalysis revealed progressive hepatic accumulation of EON-2, with liver concentrations reaching approximately 50 µg/g at day 1 and increasing to nearly 80 µg/g by day 45 (Figure 3B, right panel). To assess editing specificity, deep RNA sequencing was performed on the livers of these mice. RNA-seq-derived editing levels were consistent with dPCR measurements, confirming robust on-target quantification. No bystander editing was detected in the *B4galt1* transcript for EON-2 (Figure S2). To evaluate whether the comparatively lower peak editing efficiency observed in E3L.CETP mice versus C57BL/6J mice (∼15% versus ∼27%, respectively) reflected altered endogenous editing capacity, hepatic *Adar1* and *Adar2* transcript levels were assessed. No significant differences were detected between EON-2–treated and C57BL/6J mice, indicating that the reduced editing efficiency was not attributable to impaired expression of these genes (data not shown).

**Figure 3.**
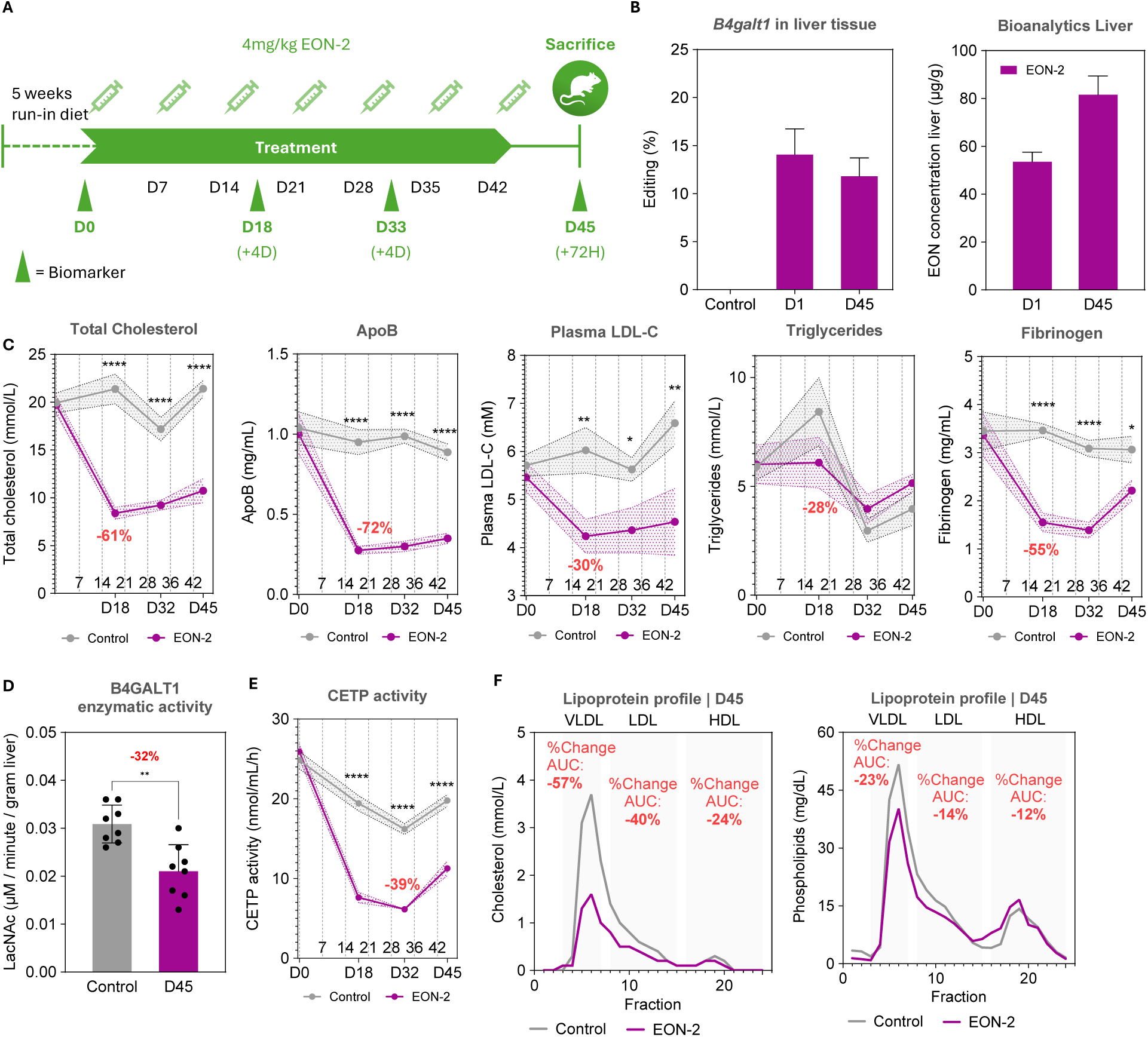
Enhanced *B4galt1* RNA editing by EON-2 drives sustained lipid modulation and optimized therapeutic outcomes in a cardiovascular disease model (EON-2–treated mice versus controls) (A) Study design: E3L.CETP mice were maintained on a Western-style diet for 5 weeks then given weekly intravenous tail-vein injections of EON-2 (4 mg/kg). (B) Proportion of *B4galt1* edited by EON-2 (left: dPCR) and hepatic accumulation of EON-2 (right: bioanalysis) at days 1 and 45. (C) Reductions in cholesterol, ApoB, LDL-cholesterol, triglycerides, and fibrinogen over 45 days. (D) B4GALT1 galactosyltransferase activity at day 1. (E) CETP activity over 45 days. (F) Cholesterol and phospholipids profiles after 45 days. \**p□* <□ 0.05 is \*\**p□* <□ 0.01 \*\*\**p* < 0.0001. Error bars represent standard deviation. ApoB, apolipoprotein B; AUC, area under the curve; BGALT1, beta-1,4-galactosyltransferase 1; CETP, cholesteryl ester transfer protein; dPCR, digital polymerase chain reaction; D, day; EON, editing oligonucleotide; HDL, high-density lipoprotein; LDL-C, low-density lipoprotein cholesterol; VLDL, very low-density lipoprotein.

The robust editing efficiency translated to pronounced lipid-lowering effects. By day 18, reductions were observed in mean total cholesterol (−61%; *p* < 0.0001), ApoB (−72%; *p* < 0.0001), plasma LDL-C (−30%; *p* < 0.01), triglycerides (−28%; *p* > 0.05), and fibrinogen (−55%; *p* < 0.0001) (Figure 3C). Reductions in cholesterol, ApoB, plasma LDL-C, and fibrinogen were sustained throughout the study. Notably, EON-2 treatment successfully prevented the acute triglyceride peak typically observed in E3L.CETP mice during the first 14–21 days of Western diet feeding. Following this period, both the EON-2 and control groups exhibited a similar downward trend toward a metabolic plateau. This convergence may be due to high inter-individual variability and potential physiologic adaptation of the control group to chronic dietary stress, in which the initial hyperlipidemic surge is followed by stabilization of lipid clearance and secretion. Fibrinogen levels demonstrated a mild rebound towards the end of the study, although its concentration remained approximately 30% lower than that of vehicle-treated controls, indicating that accelerated, editing-driven clearance still predominated over compensatory hepatic overproduction. Importantly, although *B4galt1* mRNA and B4galt1 protein levels remained unchanged, EON-2 treatment resulted in a statistically significant (32%) reduction in hepatic B4galt1 galactosyltransferase activity compared with control livers (*p* < 0.01; Figure 3D), confirming functional modulation of the enzyme. Furthermore, significant suppression of cholesteryl ester transfer protein (CETP) activity (a key mediator of lipoprotein remodeling) that transfers cholesteryl esters (CE) from high-density lipoprotein (HDL) to ApoB-containing lipoproteins in exchange for triglycerides showed a 39% decrease by day 32. This was maintained throughout the study (*p* < 0.0001 across multiple time points), providing further evidence of the potent metabolic impact of RNA editing to further attenuate pro-atherogenic lipid transfer activity (Figure 3E).

Pooled plasma lipid profiling on day 45 revealed significant alterations across cholesterol fractions, with substantial reductions in very low-density lipoprotein (VLDL) cholesterol (−57%), LDL-C (−40%), and HDL cholesterol (−24%) compared with control mice (Figure 3F). Similarly, phospholipid content within lipoprotein fractions was consistently reduced (VLDL phospholipids by −23%, LDL phospholipids by −14%, and HDL phospholipids by −12%), aligning closely with observed cholesterol reductions. This suggests a comprehensive remodeling of lipoprotein composition, potentially promoting reverse cholesterol transport pathways mediated by HDL.

### Transcriptomic and proteomic responses reveal B4galt1 RNA editing-driven metabolic adaptations

To elucidate the molecular mechanisms underlying the systemic metabolic adaptations induced by *B4galt1* RNA editing, we performed integrated transcriptomic and proteomic analyses of liver and plasma tissues on day 45 following EON-2 treatment of E3L.CETP mice, specifically focusing on the biological pathways modulated by *B4galt1* RNA editing. This multi-omics analysis aimed to capture secondary and tertiary network responses beyond direct editing targets, mapping systemic consequences of glycosylation modulation across lipid, coagulation, and homeostatic programs.

Pathway-level enrichment analysis in the liver revealed consistent downregulation of lipid-associated processes, including lipid/fatty acid metabolism (blue), glucose/broader metabolic regulation (orange), and sterol/cholesterol or regulatory processes (grey) (quadrant Q3, Figure 4A) across mRNA and protein levels. In contrast, the lipid export pathway (blue) was upregulated at both transcript and protein levels (Q1), suggesting a compensatory response aimed at facilitating lipid clearance despite the overall suppression of lipid metabolic flux. Modest upregulation was observed in glycosylation/carbohydrate pathways (pink, Q2), suggesting compensatory metabolic reprogramming in response to reduced lipid flux (Figure 4A). Key regulators of cholesterol and triglycerides homeostasis and uptake, such as ApoB, ApoE, ApoC1, and ApoC3, were downregulated at both RNA and protein levels, while fibrinogen-related genes (*Fga*, *Fgb*, *Fgg*) also showed reduced expression, corroborating the lowered fibrinogen levels (Figure 4B).

**Figure 4.**
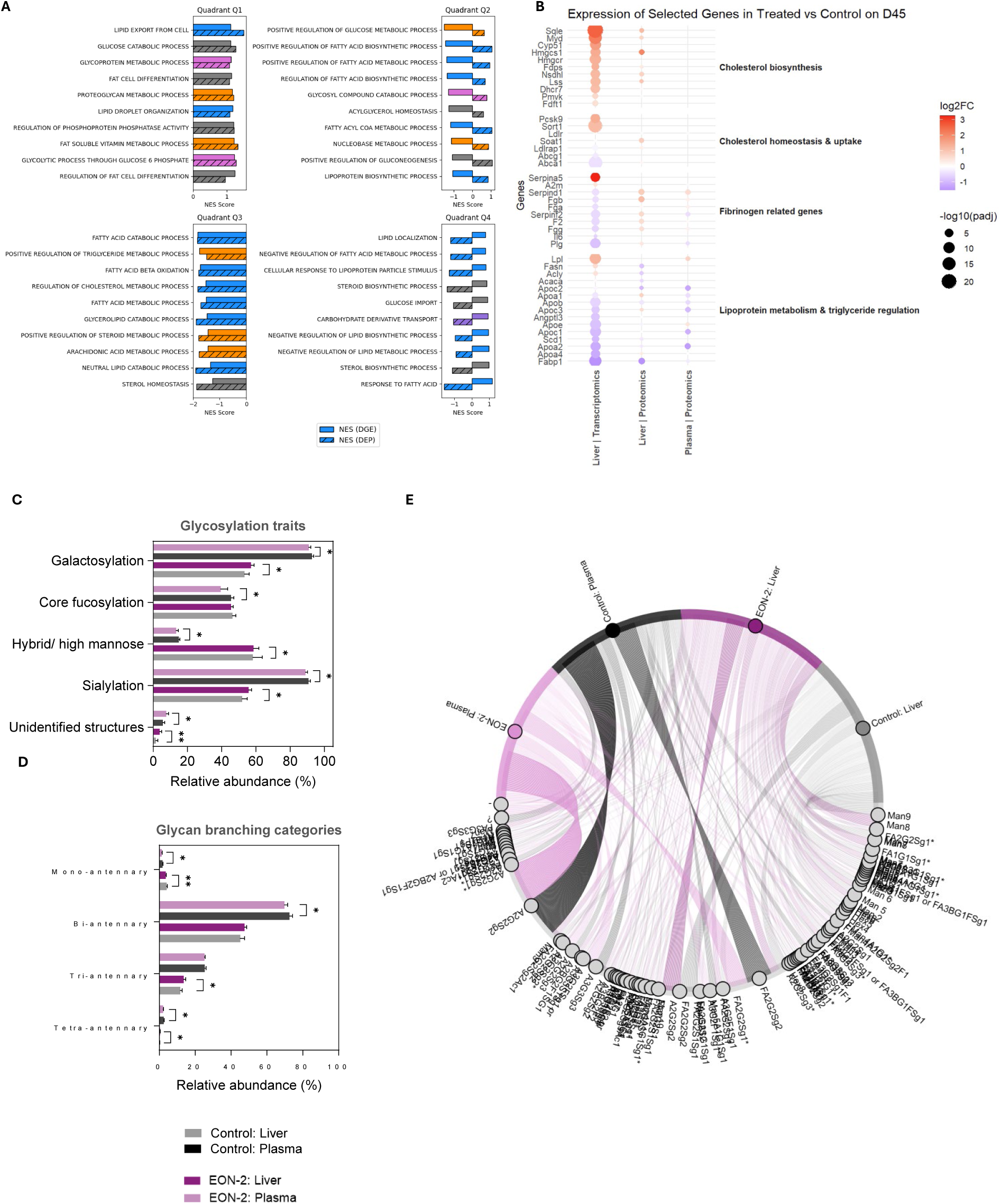
Transcriptomic and proteomic responses reveal EON-2–mediated *B4galt1* RNA editing–driven metabolic adaptations in treated E3L.CETP mice versus controls (Day 45) (A) Pathway-level enrichment analysis in liver tissue. Quadrants indicate the directionality of pathway regulation across mRNA and protein layers (Q1: transcript up/protein down; Q2: transcript up/protein up; Q3: transcript down/protein down; Q4: transcript down/protein up). Solid bar: normalized enrichment score (NES) from differential gene expression; Hatched bar: NES from differential expressed protein. Blue: Lipid/fatty acid metabolism; Orange: Glucose/broader metabolic regulation; Purple: Carbohydrate transport; Grey: Sterol/cholesterol or regulatory processes; Pink: Carbohydrate metabolism & glyco-processing. (B) Expression levels of key regulatory genes and proteins involved in cholesterol biosynthesis and homeostasis, lipid metabolism and triglyceride regulation, and fibrinogen. (C–E) Glycomic profiling of liver tissue and plasma. Error bars indicate standard deviation. \**p* □<□ 0.05 \*\**p□* <□ 0.01. BGALT1, beta-1,4-galactosyltransferase 1; EON, editing oligonucleotide; NES, normalized enrichment score.

Interestingly, within the cholesterol biosynthesis module, genes including *Sqle*, *Cyp51*, and *Hmgcr* were transcriptionally upregulated. However, this RNA-level activation did not translate to increased protein expression, suggesting post-transcriptional regulation or translational buffering that limits biosynthetic output despite transcriptional activation (Figure 4B).

To capture B4galt1-dependent alterations in protein glycosylation, comprehensive glycomic profiling was performed. This analysis revealed distinct, tissue-specific changes in glycosylation patterns in response to EON-2 treatment. In plasma, EON-2 treatment induces coordinated reductions in galactosylation, sialylation, and core fucosylation compared with control. As most circulating glycoproteins are synthesized and secreted by hepatocytes, these plasma glycomic changes likely reflect altered hepatic glycosylation prior to systemic release. In contrast, hepatic glycosylation profiles showed relative stability or modest increases across several traits following EON-2 treatment (Figure 4C). Analysis of glycan branching categories revealed that bi-antennary glycans remained the predominant structures across tissues, with higher-order branching contributing only marginally to the overall glycome (Figure 4D). At the glycan species level, EON-2 selectively reduced complex structures such as FA2G2Sg2 and A2G2Sg2 in plasma (Figure 4E).

### Temporal proteomic profiling in plasma reveals distinct phases of metabolic, immune, and coagulation responses to B4galt1 RNA editing

To characterize the temporal dynamics of protein regulation, we performed longitudinal plasma proteomics at days 0, 18, and 45. Proteins were clustered into expression trajectories, such as Up-Down-Down, Up-Down-Up, and Up-Up-Down, based on changes across the three time points (days 0, 18, and 45) relative to the control group (Figure 5A). These patterns were quantified and visualized (Figure 5B), revealing sustained directional shifts in a substantial subset of proteins by day 45.

**Figure 5.**
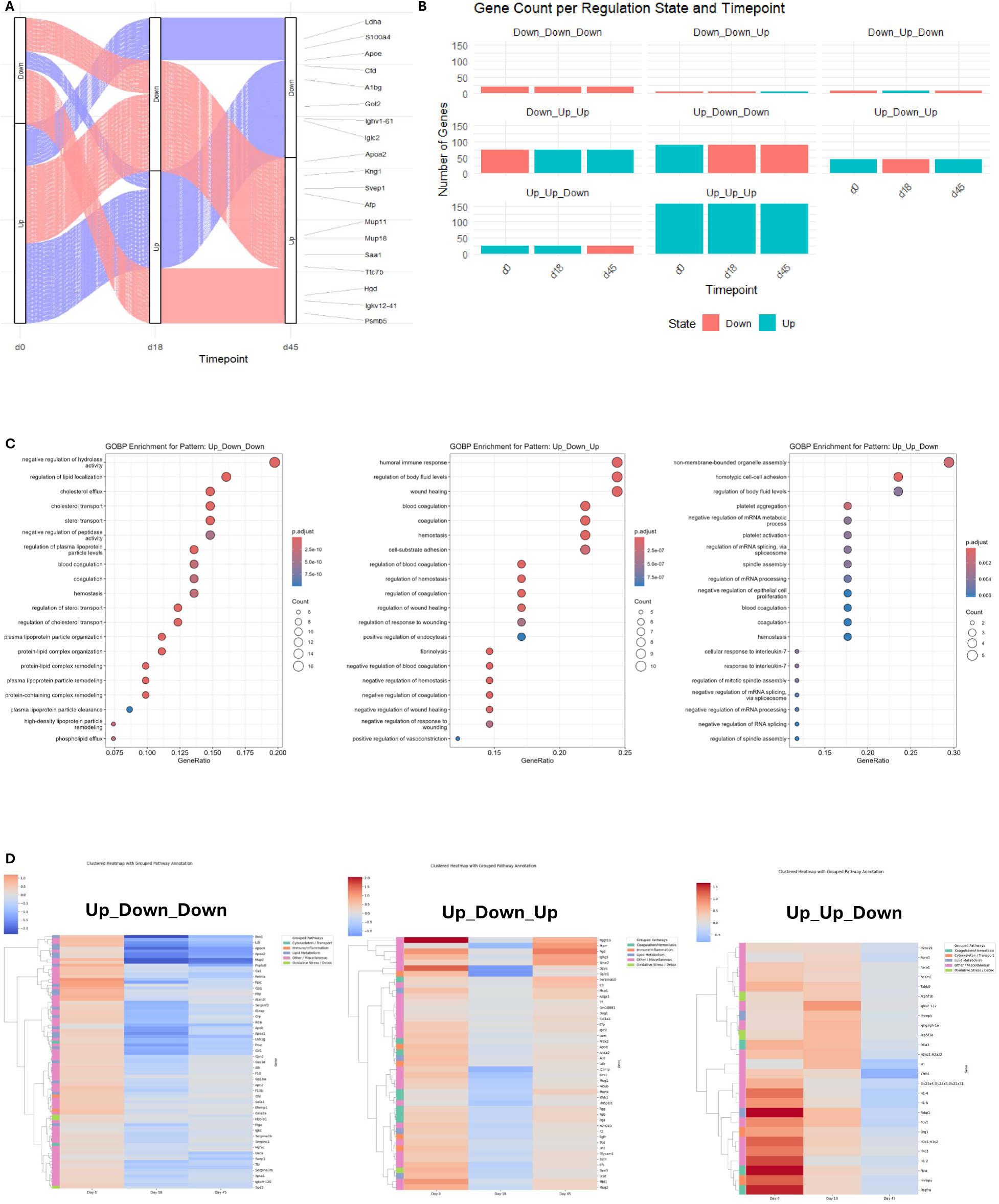
Temporal proteomic profiling reveals distinct phases of metabolic, immune, and coagulation responses to *B4galt1* EON-2 RNA editing. (A) Longitudinal plasma proteomics at days 0, 18, and 45. (B) Gene count per regulation state and time point. C) Gene ontology biological process enrichment by regulation pattern cluster (Left: Up-Down-Down; Middle: Up-Down-Up; Right: Up-Up-Down). (D) Gene-level dissection by regulation pattern cluster (Left: Up-Down-Down; Middle: Up-Down-Up; Right: Up-Up-Down). GOBP, gene ontology biological process.

The Up-Down-Down group, representing proteins transiently induced on day 18 and suppressed thereafter, was enriched in lipid regulatory pathways, including cholesterol efflux, lipoprotein remodeling, and plasma lipoprotein particle assembly (Figure 5C, left panel). This reflects an early activation followed by sustained repression, aligning with persistent reductions in circulating lipids. The Up-Down-Up trajectory, associated with coagulation and vascular homeostasis (e.g., blood coagulation, hemostasis regulation, complement activation), likely reflects feedback or compensatory responses to reduced fibrinogen abundance (Figure 5C, middle panel). The Up-Up-Down group encompassed inflammation and immune-related proteins (e.g., cytokine production, neutrophil-mediated immunity), suggesting delayed suppression of early inflammatory responses potentially triggered by RNA editing or oligonucleotide sensing (Figure 5C, right panel).

Gene-level dissection further revealed early upregulation and sustained suppression of *Apoc4*, *Apoa2, Apob, Apoa1, Pnpla8 and Sod3* in the Up-Down-Down cluster, implicating progressive inhibition of lipid mobilization and redox modulation (Figure 5D, left panel). The Up-Down-Up group included key coagulation and matrix remodeling genes (*Fgg, Fgb, Fga, C3, Col1a1*), which showed dynamic regulation suggestive of transient suppression and later reactivation of fibrin formation, complement signaling, and extracellular matrix remodeling (Figure 5D, middle panel). In the Up-Up-Down cluster, transiently elevated genes such as *H2bc21, Fabp1, and Slc25a4/5* linked to acute-phase response and innate immunity, suggested early immune activation resolving over time (Figure 5D, right panel).

Collectively, these temporally resolved analyses highlight the capacity of *B4GALT1* RNA editing to orchestrate a multi-phased, adaptive biological response involving early lipid and immune modulation followed by sustained metabolic reprogramming.

## Discussion

This study demonstrates that *B4GALT1* RNA editing constitutes a mechanistically precise therapeutic strategy capable of reprogramming glycosylation-dependent pathways to produce metabolic corrections. We demonstrate that targeted editing of *B4galt1* RNA to produce the B4GALT1 p.Asn352Ser variant can induce multiple and sustained metabolic improvements in lipid and coagulation pathways, with editing levels ranging from ∼4% to 15%. The substantial metabolic effects observed at these modest editing efficiencies likely reflect the catalytic role of B4GALT1, whereby partial modulation of enzyme activity can influence the glycosylation and clearance of multiple circulating proteins. Notably, editing of a subset of transcripts resulted in an approximately 32% reduction in galactosyltransferase activity, consistent with functional amplification beyond the measured RNA editing percentage. Given the broad impact of glycosylation on lipoprotein stability, receptor interactions, and coagulation factor clearance, partial modulation of B4GALT1 activity can propagate into substantial systemic metabolic effects.

The difference in editing efficiency between wild-type C57BL/6J mice (27%) and E3L.CETP mice (15%) likely stems from the complexities of the hyperlipidemic model. The expanded endogenous lipoprotein pool under Western diet conditions may compete with LNPs for hepatic uptake, limiting intracellular EON concentration. While the wild-type mouse study captured peak pharmacodynamic effects, the E3L.CETP data reflect steady-state editing during chronic (45-day) administration. Additionally, metabolic adaptation to the Western diet could alter *B4galt1* mRNA processing or transport, reducing the residence time of target mRNA in the nucleus and/or limiting the availability of the mRNA for ADAR-mediated deamination. Importantly, the lower efficiency appears not to be due to impaired expression of *Adar1* or *Adar2,* as these transcript levels remained unchanged in the livers of EON-treated E3L.CETP mice compared with wild-type C57BL/6J mice. However, it is also possible that changes in signaling pathways regulating editing activity may result in differential post-translational modifications of ADAR1 or ADAR2 in the E3L.CETP mice.

Using EONs to recruit endogenous ADAR enzymes, we achieved precise and tunable modulation of *B4galt1* mRNA, leading to reductions in total cholesterol, triglycerides, ApoB, LDL-C, CETP, and fibrinogen, all without altering *B4galt1* mRNA or protein expression levels. These molecular and phenotypic effects were supported by transcriptomic and proteomic remodelingNotably, partial RNA editing resulted in an approximately 32% reduction in galactosyltransferase activity, demonstrating that editing a subset of transcripts can lead to a disproportionately larger decrease in overall enzymatic function.

Multi-omics analysis revealed a dynamic hepatic response, with transient suppression of lipid metabolism and sustained downregulation of cholesterol biosynthesis pathways. These findings suggest that initial changes in glycoprotein processing may trigger broader adjustments in hepatic lipid metabolism over time.

Notably, the plasma glycomic remodeling observed in our study recapitulates glycosylation phenotypes observed in carriers of hypomorphic B4GALT1 variants, supporting a shared mechanism in which partial reduction of B4GALT1 activity preferentially affects terminal galactosylation and sialylation of circulating glycoproteins. The relative stability of hepatic glycosylation suggests the presence of compensatory, wave-like remodeling processes that mitigate broader disruption of glycan architecture. For example, downregulation of *CYP8B1* was observed, which encodes the *sterol 12-alpha-hydroxylase enzyme*. *This enzyme* is a key regulator of bile acid composition, determining the ratio of cholic acid to chenodeoxycholic acid. Reduced *CYP8B1* expression shifts the composition of bile acids toward an increased ratio of chenodeoxycholic acid, which is an agonist of the farnesoid X receptor (FXR). FXR activation has been shown to suppress bile acid synthesis and reduce the conversion of cholesterol into bile acids, which may result in accumulation of cholesterol in the liver and consequently down-regulation of the cholesterol synthetic pathway. However, given the complex and context-dependent effects of FXR signaling on cholesterol and bile acid homeostasis, the functional consequences of *CYP8B1* downregulation in this setting require further investigation.

Our findings were validated in the E3L.CETP mouse model, a well-established and human-relevant model for studying atherogenic dyslipidemia and CVD.^18, 20, 21^ Under prolonged Western dietary stress, *B4galt1* editing effectively attenuated plasma lipids and lipoproteins and suppressed CETP activity comparable with, or exceeding, the effects reported with lipid-lowering therapies such as proprotein convertase subtilisin/kexin type 9 (PCSK9) and CETP inhibitors.^19, 20, 22–25^ Importantly, unlike approaches that silence or ablate gene function (e.g., siRNA or monoclonal antibodies), RNA editing preserved *B4galt1* expression, allowing physiologic glycosylation to be maintained while selectively modifying enzyme activity.^13^ The resulting biochemical profile underscores a mechanistically targeted post-transcriptional strategy that reprograms lipid metabolism, dampens CETP activity, and remodels lipoprotein architecture, while the consistent hepatic uptake and availability of EON-2 confirm effective liver targeting. Temporal plasma proteomics revealed distinct phased responses to *B4galt1* RNA editing, characterized by early lipid regulatory changes, dynamic modulation of coagulation pathways, and transient immune activation. These adaptive shifts align with sustained metabolic reprogramming and highlight the coordinated systemic response following treatment. Collectively, these results demonstrate that *B4GALT1* RNA editing is a promising and potent therapeutic strategy for the comprehensive long-term management of CVD.

The relevance of subtle targeting of B4GALT1 is further underscored by human genetic evidence. Complete loss-of-function mutations in *B4GALT1* cause a congenital disorder of glycosylation, B4GALT1-cogenital disorder (CDG), characterized by multisystem defects.^26, 27^ Notably, patients with either homozygous or heterozygous B4GALT1-CDG exhibit marked reductions in ApoB and LDL-C, with a 26% decrease in CETP activity, mirroring the biomarker profile observed in our EON-treated animals.^26^ These parallels validate the causal role of B4GALT1 in regulating lipoprotein clearance and support the feasibility of therapeutically replicating protective alleles in a controlled non-pathogenic manner via targeted RNA editing. Importantly, the therapeutic window for B4GALT1 modulation is intrinsically narrow: although the ∼50% reduction in enzymatic activity observed in human p.Asn352Ser carriers is strongly cardioprotective, complete B4GALT1 deficiency is detrimental in humans and lethal or developmentally catastrophic in mice. By achieving 15–27% RNA editing at the 4 mg/kg dose, we elicited robust metabolic improvements while remaining well below the threshold associated with these adverse phenotypes.

The divergent fibrinogen trajectories across studies are, in part, timing-dependent: measuring the EON-2 group only 24 hours after the last LNP dose captures the acute-phase surge in hepatic fibrinogen synthesis, whereas the ≥10-day interval used for EON-2 allows that surge to resolve, unmasking the accelerated glycosylation-dependent clearance that dominates once hepatic editing exceeds ∼10%.

This work introduces a new paradigm in genetic medicine; one that expands the focus from correcting pathogenic mutations to engineering evolutionarily advantageous variants, thereby enhancing disease resistance.^12^ The ability to fine tune enzymatic output without altering gene expression offers a novel therapeutic approach, enabling controlled modulation of protein activity while preserving physiologic regulation.

Nevertheless, our study has limitations. Although the 5-week Western diet run-in period was sufficient to establish a stable hyperlipidemic phenotype in E3L.CETP mice, the overall study duration was not designed to evaluate atherosclerotic plaque development or long-term vascular outcomes. Our analyses focused on metabolic, lipid, and coagulation endpoints rather than lesion formation. Future longer-term studies will be required to determine whether sustained *B4galt1* RNA editing translates into structural improvements in atherosclerosis. While our findings suggest sustained effects, longer-term studies are needed to confirm durability and assess potential adaptive responses. An additional conceptual advantage of RNA editing-based therapies is reversibility. The present study was designed to evaluate sustained editing under repeated dosing conditions and did not include a washout phase to assess the temporal relationship between loss of editing and normalization of lipid parameters. Dedicated washout studies will be required to directly characterize reversibility of both editing and downstream metabolic effects. Additionally, current delivery approaches based on LNP formulations are not yet optimized for chronic cardiovascular indications, and the development of GalNAc-conjugated EONs will be critical for translation to hepatocyte-targeted human therapies.^28, 29^

## Conclusions

This study opens new therapeutic avenues by demonstrating that RNA editing of *B4GALT1* can selectively modulate glycosylation pathways to achieve durable metabolic benefits. The ability to introduce protective variants by targeted RNA editing *in situ* to selectively alter enzymatic output without affecting gene integrity offers a mechanism-based model for treating not only CVD but also other metabolic and immune-mediated conditions. As a reversible, programmable, and non-immunogenic platform, ADAR-based RNA editing may be a promising tool to advance precision medicine.

Future research should evaluate long-term safety, off-target profiles, and tissue specificity of editing outcomes. Translational efforts will benefit from optimized delivery systems and validation in large animal models. Expanding the editing landscape to include regulatory and protective variants may unlock a new class of therapies that improve health by deploying targeted enzymatic modulation to introduce evolutionarily conserved mechanisms of disease resistance.

## Materials and methods

### Animal welfare

The welfare of the animals was maintained following the general principles governing the use of animals in experiments of the European Communities (Directive 2010/63/EU) and Dutch legislation (the revised Experiments on Animals Act, 2014).

### *In vivo* experimental design and dosing rationale

Two *in vivo* studies were conducted to evaluate LNP-mediated editing comprehensively and compare different molecular versions of the EONs. All mice were housed in macrolon cages (maximum of six mice per cage) during the experiment in clean-conventional animal rooms (relative humidity 50–60%, temperature ∼21°C, light cycle 7 am to 7 pm). Ear punch-holes marked individual animals. Mice were supplied with heat-sterilized tap water *ad libitum* and food as described for the different groups.

Female mice were used in all *in vivo* studies (wild-type C57BL/6J and E3L.CETP) to ensure consistency across experiments and because female E3L.CETP mice on a Western type diet develop stable, reproducible diet-induced hyperlipidemia and show reliable responses to lipid-lowering interventions.^31^ Group sizes were based on prior experience with these models and historical variability of primary lipid endpoints. The validation study used n = 8 animals per group, while exploratory pharmacokinetic/pharmacodynamic studies in wild-type mice were conducted as pilot experiments (n = 3 per time point).

Dose levels were selected based on a stepwise evaluation strategy integrating pharmacokinetic, pharmacodynamic, and *in vitro* potency data. An initial 2 mg/kg dose was used to characterize plasma and hepatic exposure in wild-type mice. A 3 mg/kg dose was subsequently evaluated for pharmacodynamic assessment of editing efficiency and biomarker modulation. For the confirmatory and disease-model studies, a 4 mg/kg dose was selected based on improved *in vitro* potency and initial *in vivo* performance, as this exposure level enhanced peak editing while providing robust downstream metabolic effects. In addition, dosing intervals were selected based on observed editing kinetics. In wild-type mice, time points were chosen to capture peak and decline phases following single administration. In the E3L.CETP study, weekly dosing was implemented to maintain sustained hepatic editing over the 45-day treatment period, consistent with the editing durability observed in exploratory studies.

### Exploratory kinetics study in wild-type mice (EON-1)

The initial study aimed to characterize the pharmacokinetics and editing efficiency of the first-generation editing oligonucleotide, EON-1. EON-1 (RM4830) is a chemically modified antisense editing oligonucleotide targeting *B4galt1/B4GALT1* mRNA (see Supplementary Methods S1 for editing oligonucleotide design and chemistry). The precise antisense oligonucleotide sequence for EON-1 is disclosed in the international patent application no. WO 2024/121373. EON-1 comprises mixed, 2’-O-MOE, 2’-O-Me and 2′-F ribose modifications with phosphorothioate/phosphodiester linkages, and was synthesized and purified using standard solid-phase phosphoramidite and RP-UPLC-MS methods. Detailed information on oligonucleotide design, chemical modifications, synthesis, purification, and analytical characterization is provided in Supplementary Methods (S1–S4). Female wild-type mice (C57BL/6J), 10–12 weeks old (n = 3 per time point), received a single intravenous 2 mg/kg dose of LNP-containing EON-1 (see Supplementary Methods S5 for LNP formulation and S6 LNP characterization). Plasma EON concentration was assessed at multiple time points post treatment (10 minutes, 2, 6, 16, 24, 48, 96, 196 hours). Hepatic EON concentrations and RNA editing were assessed at 2, 6, 16, 24, and 48 hours, as well as 4 and 7 days post administration. This pilot study provided critical insights into the temporal EON distribution and editing, guiding dose selection for subsequent studies. In addition, a confirmatory *in vivo* comparison of EON-1 and EON-2 was conducted in wild-type C57BL/6J mice under comparable dosing conditions to assess relative editing efficiency and hepatic exposure.

### Exploratory pharmacodynamics study in wild-type mice (EON-1)

This study aimed to characterize the pharmacodynamics response of the first-generation editing oligonucleotide, EON-1. The precise antisense oligonucleotide sequence for EON-1 (RM4830) is disclosed in the international patent application no. WO 2024/121373. Female wild-type mice (C57BL/6J), 10–12 weeks old (n = 3 per time point, plus a control group [n = 8]; N = 21 total), were administered a single intravenous 3 mg/kg dose of LNP-containing EON-1. RNA editing, EON liver concentration, and biomarker measurements were assessed at multiple time points (2, 4, and 7 days post administration). This pilot study provided critical insights into the temporal dynamics of editing and biomarker response, guiding biomarker measurement in the following studies.

### *In vitro* editing experiments in primary human hepatocytes

The *in vitro* editing efficiency of EONs was evaluated in PHH (BioIVT) using transfection as the delivery method. For transfection, 0.5 × 10⁵ PHH were first seeded in INVITROGRO cryoplateable medium supplemented with TORPEDO Antibiotic Mix in 96-well plates and allowed to adhere for 24 hours. Cell culture medium was first refreshed for INVITROGRO Heat-Inactivated (HI) medium and EONs were then delivered using RNAiMAX (Thermo Fisher Scientific) at a 1:3 W:V ratio, with final EON concentrations ranging from 2 nM to 225 nM. Control wells received transfection medium without EON (vehicle-only control). The transfection medium was refreshed 24 hours post transfection, and cells were lysed 48 hours after the initial EON exposure.

For dPCR analysis, cells were lysed in homogenization buffer and total RNA was extracted using the Maxwell^®^ RSC simplyRNA Cells kit (Promega), following the manufacturer’s protocol. cDNA synthesis was performed using Maxima Reverse Transcriptase, random hexamer primers, 10 mM deoxynucleoside triphosphate mix (all Thermo Fisher Scientific), and nuclease-free water (Invitrogen). dPCR was carried out using the QIAcuity 4, 5-plex system, a QIAcuity PCR kit, and 96-well 8.5K nanoplates (all Qiagen). Data analysis was conducted with QIAcuity Suite Software (Qiagen).

### Validation study in Western diet-fed E3L.CETP mice (EON-2)

Building on insights from the exploratory studies, the third study tested the second-generation EON-2 in a well-established humanized model for lipoprotein metabolism, the E3L.CETP mouse model. EON-2 is a second-generation *B4galt1/B4GALT1* RNA editing oligonucleotide (see Supplementary Methods S1 for editing oligonucleotide design and chemistry). Its sequence retains the same mixed 2′MOE/2′OMe/2′F and phosphorothioate/phosphodiester backbone chemistry as EON-1, with sequence refinements to enhance editing efficiency. Details of the underlying oligonucleotide chemistry and formulation approach are provided in Supplementary Methods S1–S6. Female E3L.CETP mice aged 8–12 weeks (n = 8 per group) were obtained from the specific pathogen free breeding stock at TNO-Metabolic Health Research (Leiden, The Netherlands) and maintained on a Western-style diet containing 0.15% cholesterol and 15% saturated fat for 5 weeks before treatment. Mice received repeated intravenous tail-vein injections of 4 mg/kg of LNP-formulated EON-2 on days 0, 7, 14, 21, 28, 35, and 42 and were sacrificed on day 45.

This validation study aimed to confirm the activity, durability, and therapeutic potential of EON-2. This regimen was designed to maximize RNA editing efficiency and assess sustained metabolic responses over a 45-day period.

### Diet

Throughout the validation study, E3L.CETP mice were maintained on a Western-type diet as described by Nishina et al. (1990).^32^ The diet contained cholesterol (0.15% w/w, final concentration) and 15% cacao butter. It was stored in vacuum-sealed bags in a temperature-controlled (–20°C) environment. Fresh diet was provided weekly to ensure consistency in lipid metabolism conditions.

### Blood and plasma

Samples were obtained after 4 hours of fasting. Animals were placed under a red lamp for at least 10 minutes. The tail was fixed and the mice could move freely during blood collection to avoid additional stress. An incision was made in the tail vein to collect tail blood using microvette tubes containing EDTA-dipotassium salt (CB 300 K2E microvettes; Sarstedt, Nürnbrecht, Germany). Tubes were placed on ice immediately. Plasma was obtained after centrifugation (10 min at 9000 x g) of the samples in a laboratory centrifuge at 4°C. Plasma (supernatant after centrifugation) was separated, stored in a 1.5 mL tube, and stored at –80°C until further analysis.

### Blood parameters

Total plasma cholesterol and triglycerides were quantified using the “Cholesterol Gen.2” and “Trigl” kits from Roche/Hitachi (Cat. No. 03039773190, Roche Diagnostics GmbH, Mannheim, Germany). Plasma LDL cholesterol was determined with the mouse LDL-C assay kit from Chrystal Chem (Cat. No. 79980, Crystal Chem Inc., Elk Grove Village, IL, USA). Plasma fibrinogen was assessed using the mouse Total Fibrinogen ELISA kit from Innovative Research (Cat. No. MFBGNKT, Innovative Research Inc., Novi, MI, USA). Plasma ApoB concentrations were determined with the mouse ApoB ELISA from Abcam (Cat. No. ab230932, Abcam, Cambridge, MA, USA). Lipoprotein profiles were generated by fast protein liquid chromatography analysis using an ÄKTA™ apparatus; samples were pooled per group, and cholesterol and phospholipids in the fractions were measured with the Cholesterol Gen.2 kit from Roche and the Phospholipids kit from Instruchemie (Cat. No. 03039773190, Roche Diagnostics GmbH, Mannheim, Germany, and Cat. No. 3009, Instruchemie B.V., Delfzijl, The Netherlands).

### Sacrifice and harvest of tissues

Mice were sacrificed following a 4-hour fast via heart puncture under isoflurane anesthesia. Sacrifice was performed with group randomization based on the time of sacrifice. Plasma, collected in lithium heparin, was obtained by heart puncture and stored at –80°C until further use. The liver was isolated and processed. The liver was weighed and divided into five portions: one piece from the medial lobe was fixed in 10% formalin and sent to the sponsor, while four additional pieces were snap-frozen in liquid nitrogen and stored individually at –80°C; one piece was designated for high-performance liquid chromatography analysis (20–60 mg), another for editing analysis by droplet dPCR (ddPCR; 20–60 mg), and the remaining two reserved for future analyses.

### RNA sequencing and bioinformatic analysis

Liver tissues were harvested, flash-frozen in liquid nitrogen, and stored at –80°C (or alternatively preserved in RNAlater® from Thermo Fisher Scientific) (Cat. No. AM7020, Thermo Fisher Scientific, Waltham, MA, USA) until RNA extraction. Total RNA was then extracted using the Qiagen RNeasy Mini Kit (Cat. No. 74104, Qiagen, Hilden, Germany) following the manufacturer’s instructions, with tissue homogenization performed using a bead homogenizer and genomic DNA removed by DNase I treatment. RNA concentration and purity were assessed by NanoDrop spectrophotometry, and RNA integrity was evaluated using an Agilent 2100 Bioanalyzer, with only samples exhibiting an RNA Integrity Number of ≥7 proceeding to further analysis. Polyadenylated RNA was enriched using the NEBNext Poly(A) mRNA Magnetic Isolation Module (Cat. No. E7490, New England Biolabs, Ipswich, MA, USA), or, alternatively, rRNA was depleted using the NEBNext rRNA Depletion Kit, as appropriate. Enriched mRNA was reverse-transcribed into first-strand complementary DNA (cDNA) using the NEBNext First Strand Synthesis Module and then converted into double-stranded cDNA through second-strand synthesis. This double-stranded cDNA was subsequently fragmented to an average size of 200 bp using the NEBNext Ultra II Directional RNA Library Prep Kit for Illumina (Cat. No. E7760, New England Biolabs, Ipswich, MA, USA), which also included end-repair, A-tailing, and adapter ligation. Libraries were then amplified with indexing primers via PCR and purified using AMPure XP beads (Cat. No. A63880, Beckman Coulter, Brea, CA, USA); their quality and concentration were confirmed using an Agilent Bioanalyzer. Final libraries were pooled at equimolar concentrations. Library preparation followed established protocols and is summarized below. Sixty million reads (2 x 150 base pairs) per sample were sequenced using the Illumina NovaSeq platform (Illumina).^33–39^ Adapter sequences were trimmed from raw sequencing reads with BBtools BB Decontamination Using Kmers (*BBDuk*).^40^ Trimmed reads were aligned to the human reference genome (GRCh38, Ensembl release 110) using hisat2.^36^ PCR duplicates were identified and removed using SAMtools markdup.^41^

Gene-level read counts were quantified with HTSeq-count.^33^ Differential gene expression analysis was performed using DESeq2 (v1.46.0) in R (v4.4.3), which applies to a negative binomial generalized linear model to identify statistically significant expression differences between conditions.

### Off-target editing analysis

Prior to variant calling, reads were processed with GATK SplitNCigarReads to handle splice junction spanning alignments.^42^ Variants were identified using BCFtools mpileup and call performed on high-confidence reads (mapping quality ≥ 60). Sites overlapping known SNPs were excluded, and variants were filtered to retain only those consistent with A-to-I editing. Off-target RNA editing was assessed by testing for statistically significant enrichment of A-to-G variants in treated samples relative to EON-treated and untreated control conditions, using Fisher’s Exact Test (*p* < 0.05).

### Data-independent acquisition (DIA) for label-free quantification

Plasma samples collected at days 0, 18, and 45 from the validation study were subjected to longitudinal proteomic profiling using DIA mass spectrometry. Liver tissues (20–50 mg) were transferred to pre-chilled tubes and homogenized at 4°C using a bead homogenizer (Cat. No. 25-010, Omni International, Kennesaw, GA, USA). Protein lysates were extracted and subjected to clean-up and digestion. Peptides were fractionated and desalted prior to liquid chromatography–mass spectrometry/mass spectrometry analysis. Samples were analyzed on a Thermo Scientific Orbitrap Exploris 480 mass spectrometer, coupled with an Evosep One LC system (Cat. No. BRE725539, Evosep, Odense, Denmark). The instrument was operated in DIA mode, using 17 variable-width windows covering the mass range of 472–1143 m/z, with 39.5 m/z overlaps. MS1 spectra were acquired at 120,000 resolution with an automatic gain control target of 1000% and a maximum injection time of 45 ms. Fragmentation was performed using higher-energy collisional dissociation at 27% normalized collision energy, and MS2 spectra were acquired at 45,000 resolution with a maximum injection time of 86 ms. Peptide and protein identification were carried out using Spectronaut direct DIA for library-free searches.

### Statistics

Depending on normality, the significance of differences between the groups was calculated either parametrically or non-parametrically using the computer program GraphPad (10.4.0, GraphPad Software, Inc, San Diego, CA, USA). A Kruskal-Wallis test for several independent samples was used for non-parametric calculations, followed by a Mann-Whitney U-test for independent samples. A one-way analysis of variance for multiple comparisons was used for parametric calculations, followed by Dunnett or Bonferroni correction. A *p*-value of < 0.05 was considered statistically significant.

## DATA AND CODE AVAILIBILITY

The precise antisense oligonucleotide sequences of EON-1 (RM4830) disclosed in this study are included in the international patent application (publication no. WO 2024/121373). The precise antisense oligonucleotide sequences of EON-2 are provided in the Supplementary Materials (Supplementary Methods S1) and are included in the international patent application. All other methodological details are described to allow replication of the study design and interpretation of results. The data underlying this article will be shared on reasonable request to the corresponding author.

## ACKNOWLEDGEMENTS

The work with E3L.CETP mice was conducted at TNO-Metabolic Health Research (Leiden, The Netherlands) and funded by ProQR Therapeutics (Leiden, The Netherlands). ProQR Therapeutics was involved in the study design of this study but had no role in data collection or data analysis. Editorial support for the preparation of this manuscript (under the guidance of the authors) was provided by Becky Vickers and Suzanne Flowers, Apothecom (UK), and funded by ProQR Therapeutics (Leiden, The Netherlands). Research and data analysis was funded by ProQR Therapeutics.

## AUTHOR CONTRIBUTIONS

F.D.C., E.C.dlR., B.K., S.dK., and G.P. contributed to the conceptualization and methodology of the study. F.D.C. and E.C.dlR. performed the investigation, formal analysis, data curation, visualization, and writing – original draft. P.dB. and S.Y-E. contributed to the methodology and investigation of the in vitro studies. E.T. contributed to resources through EON synthesis and formulation. L.S., N.C., A.M.vdH., H.M.G.P., and W.B. contributed to the methodology and supervision of the in vivo studies. A.M.vdH. and H.M.G.P. additionally contributed to the conceptualization, investigation, formal analysis, and validation of the non-omics validation study using the E3L.CETP mouse model. J.J.T. contributed to methodology through the design of the EONs. S.Y-E., B.K., S.dK., and G.J.P. contributed to data interpretation and provided critical input into the experimental design and data analysis. All authors contributed to review & editing and approved the final version of the manuscript.

## DECLARATION OF GENERATIVE AI AND AI-ASSISTED TECHNOLOGIES IN THE WRITING PROCESS

During the preparation of this work the authors used ChatGPT Enterprise in order to solely for editorial support, including checks for clarity, conciseness, and language. After using this tool/service, the authors reviewed and edited the content as needed and take full responsibility for the content of the publication.

## SUPPLEMENTAL INFORMATION

Supplemental information can be found online at XXX

---

**Figure S1.**
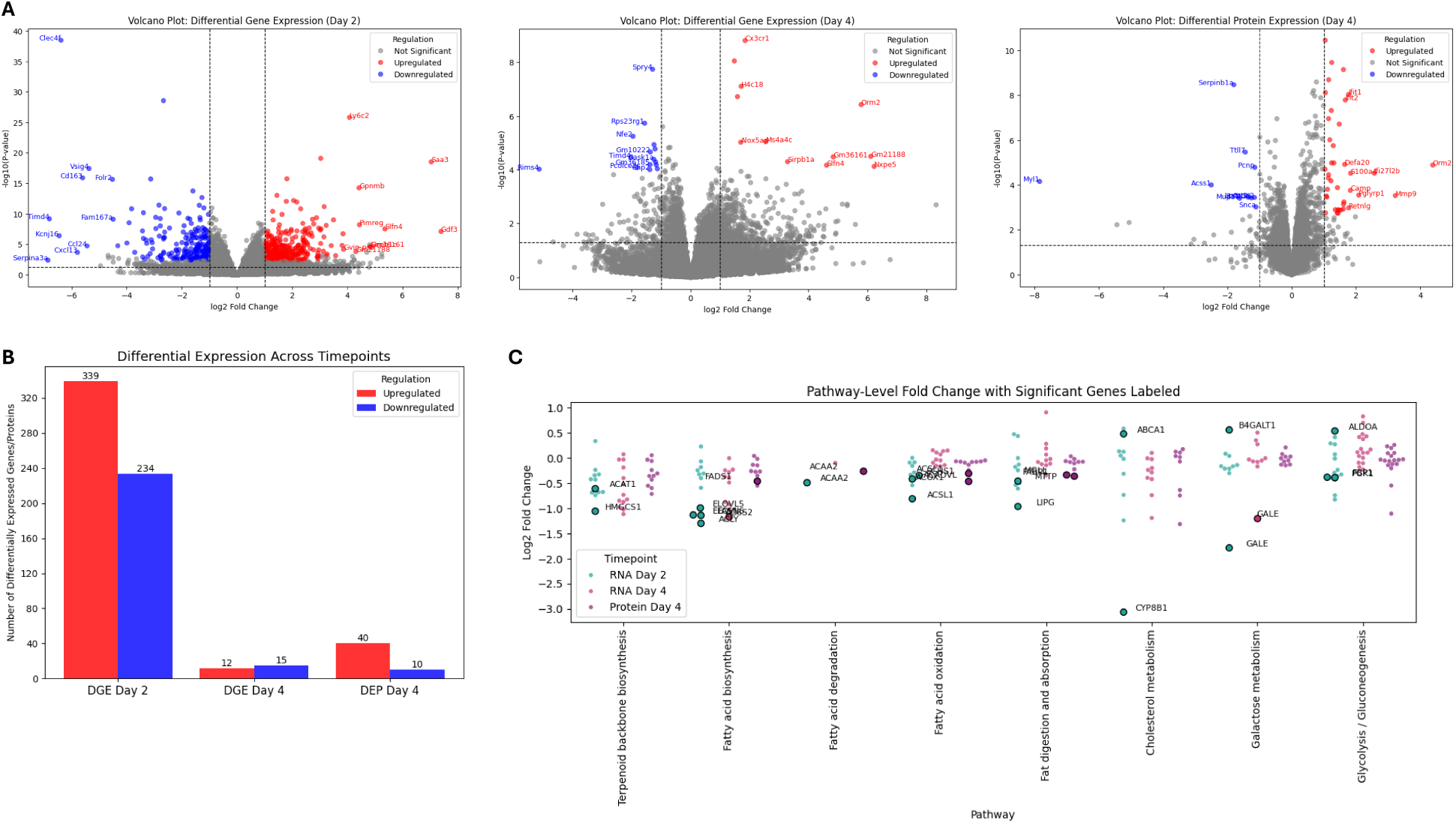
Temporal transcriptomic and proteomic changes following *B4galt1* editing with EON-1. (A−B) Transcriptomic and proteomic analyses revealed dynamic temporal regulation after *B4galt1* editing, followed by refinement of the transcriptional response by day 4. (C) Cholesterol homeostasis remained persistently downregulated at both mRNA and protein levels, driven by the downregulation of *sterol 12-alpha-hydroxylase* gene (*CYP8B1*). Labeled genes indicate those meeting statistical significance (adjusted *p*-value < 0.05). BGALT1, beta-1,4-galactosyltransferase 1; DEP, differential expressed protein; DGE, differential gene expression; EON- editing oligonucleotide; mRNA, messenger RNA;

**Figure S2.**
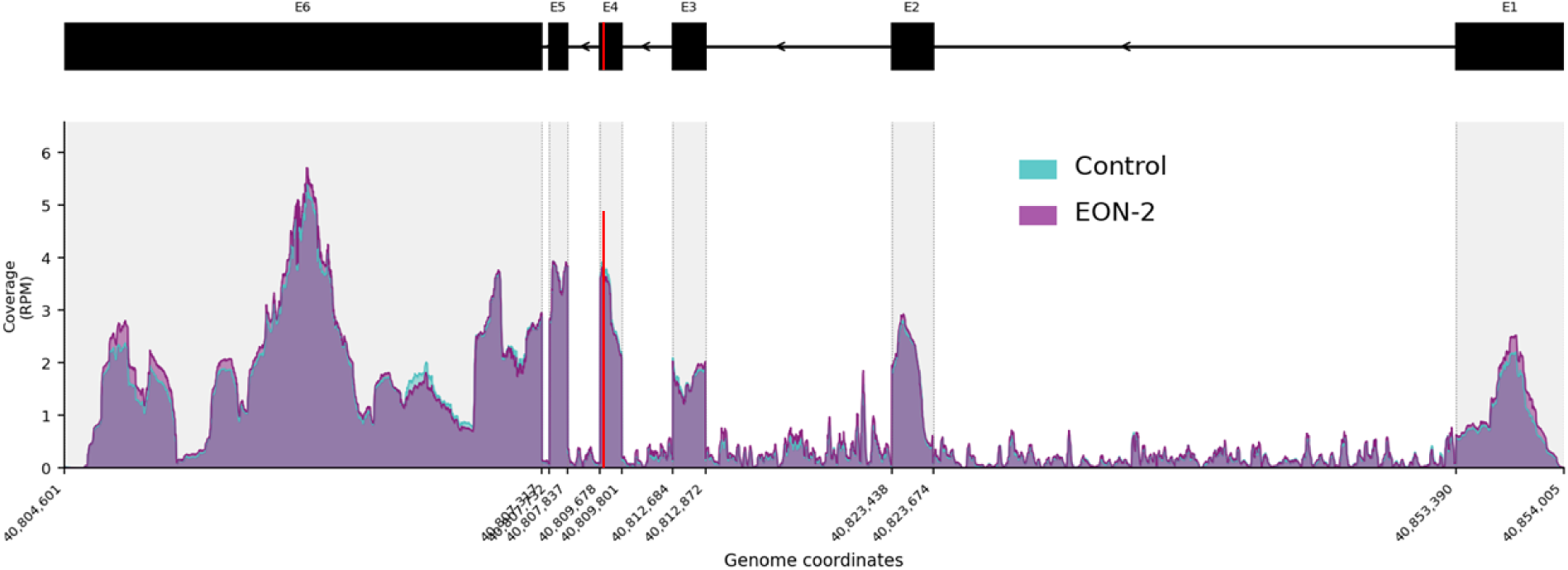
RNA-seq coverage across the *B4galt1* transcript in control and EON-2 treated samples. Per-base coverage (RPM-normalized) is plotted in genomic coordinates with control (teal) and EON-2-treated (purple) conditions overlaid for direct comparison. Introns are compressed 0.1X for visual clarity. Exon positions (E1–E6) are indicated in the gene model diagram above. Red line indicates editing site. BGALT1, beta-1,4-galactosyltransferase 1; EON- editing oligonucleotide; RPM, reads per million.

