## Supplementary Material for "ADAR RNA editing for cardiovascular disease: Targeting *B4GALT1* to modulate lipid metabolism through reduced galactosyltransferase activity"

### Supplementary Methods

#### Supplementary Methods S1. Editing oligonucleotide design and chemistry (EON-1 and EON-2)

**EON-1**

EON-1 (internal identifier RM4830) is a chemically modified antisense editing oligonucleotide designed to hybridize to murine and human *b4galt1*/*B4GALT1* mRNA and recruit endogenous ADAR enzymes to mediate site-specific A-to-I editing at the codon encoding p.Asn352. The full antisense nucleotide sequence of EON-1 is publicly disclosed in international patent application WO2024/121373A1.

EON-1 consists of a mixed-chemistry architecture incorporating:

- 2′-O-methoxyethyl (2′-O-MOE) ribose modifications
- 2′-O-methyl (2′-O-Me) ribose modifications
- 2′-fluoro (2′-F) ribose modifications
- A mixed phosphorothioate (PS) and phosphodiester (PO) internucleoside linkage backbone

The positioning of sugar and backbone chemistries was designed to balance target affinity, nuclease stability, and ADAR recruitment while maintaining favorable *in vivo* tolerability. The nucleotide opposite the target adenosine was positioned to promote selective A-to-I editing within a defined editing window.

**EON-2**

EON-2 is a second-generation *B4galt1/B4GALT1* RNA editing oligonucleotide. Its sequence retains the same mixed 2′-O-MOE/2′-O-Me/2’-F and phosphorothioate/phosphodiester backbone chemistry as EON-1, with sequence refinements to enhance editing efficiency. The full antisense nucleotide sequence of EON-2 is as follows:

m5Ce!Ge$^{\theta}$Ge$^{\theta}$m5Ue$^{\theta}$m5Ce$^{\theta}$Am*Am*Cm*Am*Am*Gm*Cf*Uf*Gf*Af*Gf*Gf*Ad*Zd*m5Ud!Gm$^{\theta}$
Gf*Ge*Uf*Uf*Cf*Af!Uf$^{\theta}$L004

m5Ce is a 2’-O-MOE modified 5-methyl cytidine, Ge is a 2’-O-MOE modified guanosine, m5Ue is a 2’-O-MOE modified 5-methyl uridine, Am is a 2’-O-Me modified adenosine, Cm is a 2’-O-Me modified cytidine, Gm is a 2’-O-Me modified guanosine, Cf is a 2’-Fluoro modified cytidine, Uf is a 2’-Fluoro modified uridine, Gf is a 2’-Fluoro modified guanosine, Af is a 2’-Fluoro modified adenosine, Ad is a deoxyadenosine, Zd is a deoxynucleotide comprising a 6-amino-5-nitro-3-yl-2(1H)-pyridone nucleobase, m5Ud is a deoxy-5-methyl uridine, * is a phosphorothioate linkage, ! is a PNdmi linkage, $^{\theta}$ is a phosphodiester linkage. L004 is GalNAc.

This sequence has been submitted in an international patent application.

**Supplementary Methods S2. Oligonucleotide synthesis**

All EONs, including EON-1, were synthesized using standard solid-phase phosphoramidite chemistry on controlled pore glass (CPG) support with commercially available protected nucleoside phosphoramidite monomers.

Following completion of chain assembly, cleavage and deprotection (ammonolysis) were performed using either aqueous ammonium hydroxide (28–30%) at 45 °C for 18 hours or gaseous ammonium hydroxide (80 psi) at 80 °C for 3 hours.

After cleavage and deprotection, crude oligonucleotide solutions were filtered to remove the solid support and washed with water to maximize product recovery.

**Supplementary Methods S3. Purification**

Oligonucleotides were purified by dimethoxytrityl (DMT)-on reversed-phase (RP) chromatography using UniPS-15 resin. After eluting DMT-off impurities and subsequent on-column DMT removal using a 6% (v/v) dichloroacetic acid (DCA) in water solution, the oligonucleotide full-length product was eluted as fractions by gradient elution with a mobile phase of 100 mM tri-ethyl ammonium acetate (TEAA) and in purified water and methanol (pH 7) (Buffer A = 2.5% (v/v) MeOH; Buffer B = 90% (v/v) MeOH). Fractions were tested for purity and pooled to the desired purity (Typically >85% purity). The product pool solution was concentrated and desalted against water for injection (or equivalent) using vivaspin ultrafiltration spin columns with 3 kDa molecular weight cut-off (or equivalent). Finally, the solution was lyophilized to obtain the oligonucleotide as a solid.

**Supplementary Methods S4. Analysis**

All oligonucleotides (i.e., crude, purification fractions, and final product) were analyzed by ion-pair reversed-phase Ultra Performance Liquid Chromatography (UPLC) coupled with single-quad mass spectrometry (IP-RP UPLC-MS) on a Waters UPLC using an ACQUITY UPLC Oligonucleotide BEH C18 Column (Waters Corporation, Milford, MA, USA; 186003949, 1.7-µm, 2.1 × 50 mm) for identity and purity determination. Typically, a 10-minute linear gradient of 10% to 40% Buffer B was used to analyze the samples where Buffer A is a solution of 385 mM HFIP (1,1,1,3,3,3-hexafluoro-2-propanol), 14.5 mM triethylamine, and 5% methanol in UPLC-grade water and Buffer B a solution of 385 mM HFIP, 14.5 mM triethylamine, and 90% methanol in UPLC grade water.

Concentration (i.e., yield) determination was performed by measuring UV absorbance at 260 nm using a UV-vis spectrometer or nanodrop system.

**Supplementary Methods S5. Lipid nanoparticle (LNP)**

EONs were encapsulated in commercial LNPs for *in vivo* delivery purposes. LNP formulations were prepared with ionizable cationic aminolipid XL-10, as described in Geisler et al. (2023).[30] The EON stock solution was prepared in phosphate-buffered saline (PBS) (1 mM $KH_2PO_4$, 155 mM NaCl, and 3 mM $Na_2HPO_4$ $0.7H_2O$, pH 7.4). The final LNP formulation was sterile, filtered over 0.22 µm sterile filter. Due to the analytical challenges in quantifying effective RNA cargo, where nominal concentrations often overestimate true loading and particle heterogeneity is high, we utilized an empirical in vivo dose-response range (2–4 mg/kg). This approach accounts for the fact that nominal dose delivery does not always correlate linearly with hepatic target engagement.

**Supplementary Methods S6. Characterization of EON LNPs**

The Zetasizer Ultra (Malvern, Worcestershire, UK) dynamic light scattering instrument was employed to determine the hydrodynamic diameter and polydispersity index of LNPs diluted in PBS. The encapsulation efficiency of LNPs and concentration were determined using the Quant-iT RiboGreen RNA assay kit (Thermo Fisher Scientific, Framingham, MA, USA).

### Supplementary Figures

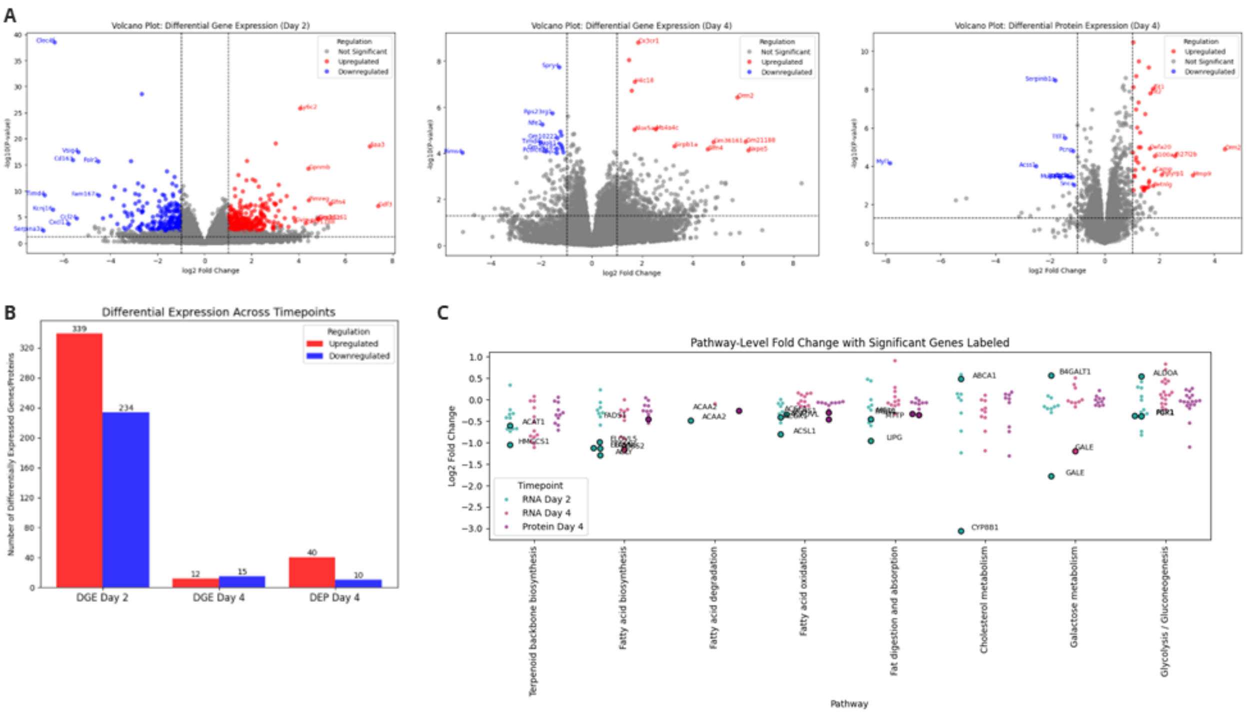


**Figure S1. Temporal transcriptomic and proteomic changes following *B4galt1* editing with EON-1**

(A−B) Transcriptomic and proteomic analyses revealed dynamic temporal regulation following *B4galt1* editing, followed by refinement of the transcriptional response by day 4. (C) Cholesterol homeostasis remained persistently downregulated at both mRNA and protein levels, driven by the downregulation of *sterol 12-alpha-hydroxylase* gene (*CYP8B1*). Labeled genes indicate those meeting statistical significance (adjusted p-value < 0.05).

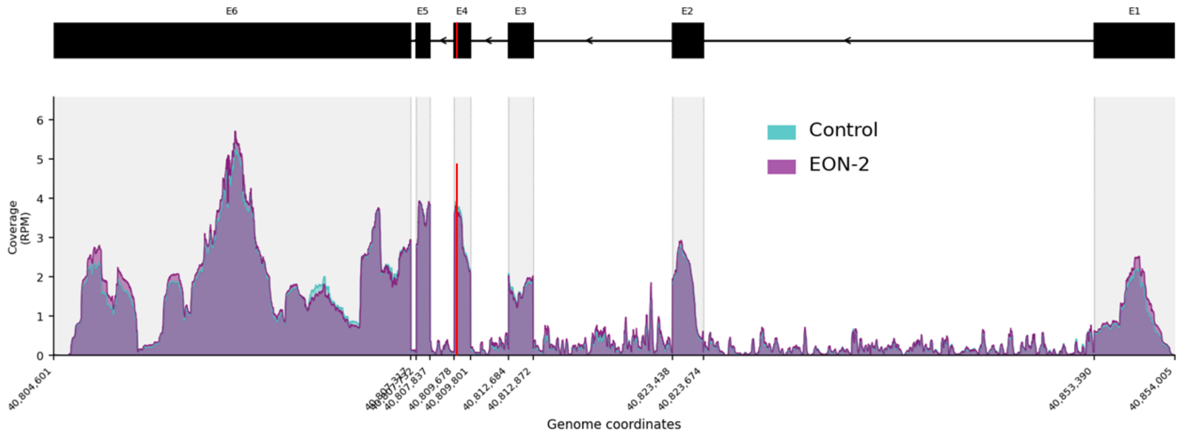


**Figure S2. RNA-seq coverage across the *B4galt1* transcript in control and EON-2 treated samples.** Per-base coverage (RPM-normalized) is plotted in genomic coordinates with control (teal) and EON-2-treated (purple) conditions overlaid for direct comparison. Introns are compressed 0.1X for visual clarity. Exon positions (E1–E6) are indicated in the gene model diagram above. Red line indicates editing site.
